# Detection of stress functional responses in bacterial populations under dry soil conditions show potential microbial mechanisms to resist drought conditions

**DOI:** 10.1101/2020.09.30.320879

**Authors:** Soumyadev Sarkar, Kaitlyn Ward, Janet K. Jansson, Sonny T.M. Lee

## Abstract

Climate change is predicted to have a negative effect on the grasslands of the United States and will be detrimental to the economy and environment. The changing precipitation levels would also have an effect on the structural and functional potential of associated soil microbiome communities, which in turn will regulate the health of the plants during stressful conditions. In this study, we applied metagenomics analyses to capture the responses of the bacterial populations under drier soil conditions. We collected soil from two sites (dry and wet) at the Konza Long-Term Ecological Research field station in Kansas, which had characteristic features of the native prairies. Soil drying resulted in a significant shift in the bacterial population at the community level. Following that, fifteen bacterial genomes were short-listed based on the availability in the public database, higher relative abundance in dry soils than in wet, and also according to their contributions in drier soil. The potential microbial mechanisms were elucidated when an in-depth analysis of the functional genes was performed. Translation elongation factor EF-Tu, thiamine biosynthesis protein, and catalase were identified as a part of the overall stress functional responses in the bacterial population in this study. We speculate that these identified bacterial populations are important for maintaining the health of the soil under dry conditions. Genes and/or pathways found in this study provide insights into microbial mechanisms that these bacterial populations might employ to resist challenging drought conditions.

## 1. Introduction

Climate change is an incessant global concern that often results in altered precipitation levels, seasonal fluctuations, warming ocean floors, etc. (VijayaVenkataRaman, Iniyan, and Goic 2012). Events like altered rainfall have a direct negative effect on agriculture and productivity (Gornall et al. 2010). In the United States, future irregularity in rainfall is predicted in the south-central, south-western, and in the northern and central grasslands of North America (Cook, Seager, and Miller 2011; Intergovernmental Panel on Climate Change 2014; Taylor et al. 2012). The increased rainfall irregularity in the mesic grasslands is associated with a reduction in soil respiration (Schimel 2018), reduced aboveground net primary productivity (ANPP) (Knapp et al. 2002; Heisler-White et al. 2009), decrease in carbon cycling processes (Heisler-White et al. 2009), altered nitrogen cycling processes (Schimel 2018), and even in decline in the level of photosynthesis (Jones et al. 2016). There are also reports of change in genotypic diversity in the grasslands as a consequence of rainfall alterations (Jones et al. 2016; Avolio, Beaulieu, and Smith 2013).

These modifications happen not only in the host but also in the associated microbiome as well (Chowdhury et al. 2019). Under changing precipitation profiles, microbiomes exhibit osmotic adjustments, optimum resource efficiency, enhanced microenvironment, stable dormancy to evade stress (Schimel 2018). It is speculated that the plant host and -associated microbiome remain complementary towards each other in this changing scenario (Lau and Lennon 2012). The shift in the microbiome has a definite impact on the plant fitness under dry conditions, and have contributed towards increasing the health of the plants during drought (Xu and Coleman-Derr 2019).

The cellular and functional responses by bacteria populations during drought conditions include induction of dormancy (forming spores) (Barnard, Osborne, and Firestone 2013), production of exopolysaccharides (Roberson and Firestone 1992), performing osmotic adjustments (Chowdhury et al. 2019), synthesis of enzymes that are extracellular in nature to sequester solutes (Chowdhury et al. 2019; Barnard, Osborne, and Firestone 2013). During drought conditions, *Actinobacteria* and *Planctomycetes* use pathways related to osmotic stress responses (Bouskill et al. 2016). Exopolysaccharides are produced in *Pseudomonas sp*. and *Acidobacteria* to resist desiccation (Roberson and Firestone 1992; Chang et al. 2007; Ward et al. 2009). *Bacillus spp*. and *Actinobacteria* possess the ability to form spores in desert-like conditions (Marasco et al. 2012).

Until now, investigations to understand soil microbiome processes under dry conditions have focussed on a more holistic approach to identifying diverse microbial populations without extracting much information about the underlying mechanisms. Although there are studies that attempt to identify the shifts in soil microbial functions under stressful conditions, there is still a clear knowledge gap that exists on how these mechanisms can be associated with the participating bacterial population during drought (Riah-Anglet et al. 2015; D. Liu et al. 2019). This study aims to fill this void, and we used metagenomic approaches at the (a) community level (b) genome level (c) functional genes level to link the impact of soil drying with mechanisms that the microbial communities adopt to resist such extreme conditions.

We collected soil samples with distinct moisture characteristics (wet and dry) from two locations in Konza Long-Term Ecological Research field station (Kansas), to investigate how changes in soil moisture levels would induce a shift in the bacterial population and in the participating metabolic genes and pathways. Our study yielded detailed information on the bacterial community, genome, and gene level, and we believe this lays the framework for other similar scientific explorations.

## 2. Materials and Methods

### 2.1. Sample sites and collection

In this study, soils were collected from two sites in Konza Long-Term Ecological Research field station in Kansas. Soil from site A (39°06 =11 N, 96°36 =48 W, 339 meters above sea level) had a higher water content of 37%, whereas site B (39°04 =39 N, 96°36 =29 W, 413 meters above sea level) soil had a lower water content of 18%. Both the sampling locations had C4 perennial grasses, resembling the characteristics of native prairies, and were not disturbed by agricultural practices.

Soil samples (n = 12) were collected at a depth of 15 cm from three locations which were minimum 10 m apart, from both sites. After removing rocks and roots, the soils were homogenized and then collected in single Ziploc bags for each site. Soils were immediately frozen in liquid nitrogen, and shipped. After receiving at the Pacific Northwest National Laboratory (PNNL), Richland, WA, the soil samples were sieved, aliquoted in 50 ml conical centrifuge tubes and stored at −80°C until further processing.

### 2.2. Shotgun metagenomics and reference-based metagenomics

DNA extraction was carried out as previously described (Chowdhury et al. 2019). Sequencing was performed on the HiSeq 2500 platform (Illumina, San Diego, CA) which generated 250-bp paired-end reads (~10 million reads per sample) from three runs. We used Kaiju ver 1.7.2 (Menzel, Ng, and Krogh 2016) to perform taxonomic classification of the high-throughput reads from the metagenomes. Clustering algorithms were executed with a 99% similarity cut off, and each sequencing read was assigned a taxon against the RefSeq database. Chimeras and sequences corresponding to plants/vertebrates were also filtered.

We short-listed and downloaded 15 genomes (~5 million bp) from the National Center for Biotechnology Information database (Sayers et al. 2020) (Supplementary Table S1). We chose these fifteen bacterial genomes based on the following criteria: 1) the bacterial populations’ higher mean relative abundance in the dry soil indicated by Kaiju analysis at the species level 2) based on their availability of the genomes in the database, and 3) contribution to the dissimilarity as analyzed from SIMPER analysis of the whole community at the species level. We downloaded the following genomes - *Azosprillum brasilense, Azospirillum thiophilum, Azotobacter chroococcum, Azotobacter vinelandii, Candidatus Nitrospira inopinata, Candidatus Nitrospira defluvii, Thiobacillus denitrificans, Nitrosococcus halophilus, Nitrosococcus watsonii, Nitrosococcus oceani, Paracoccus denitrificans, Nitrospira moscoviensis, Frankia casuarinae, Frankia inefficax*, and *Frankia Datisca glomerata*.

We mapped short-reads from each of the 12 metagenomes onto the 15 downloaded bacterial genomes using Bowtie2 ver 2.3.5.1 (Langmead and Salzberg 2012), and stored the recruited reads using samtools (Li et al. 2009). We then used anvi’o ver 6.1 (Eren et al. 2015) to process the BAM files and generate profile databases containing coverage and detection statistics of each downloaded genomes in the metagenomes.

### 2.3. Statistical analysis

We used primer v7 (Clarke and Gorley 2015) to perform all statistical analyses. For community and genome analyses, we standardized and square-root transformed (Christian, Kaestli, and Gibb 2017) the data, and tested for differences in the bacterial community using PERMANOVA analysis on Bray-Curtis similarity. SIMPER statistical analysis (cut-off: 70%) was also performed during community, genome, and functional genes analyses. We used ggplot2 in the R tidyverse package to visualize and create all figures (Gómez-Rubio 2017).

## 3. Results and discussions

To understand the soil bacterial membership and function between two soil types with distinct precipitation levels, we sampled two locations that had a defined soil moisture content difference (soil A: 37% and soil B: 18%) (Chowdhury et al. 2019). We categorized soil A as wet soil, and soil B as dry throughout this report. Our study added to the current knowledge on soil microbial functional shifts during stressful conditions. In this study, we have focused on the functional contributions, and our findings are based not only on the basis of the relative abundance but also on contributions of each member(s) at each level, i.e. community, genomes and genes. Our findings support that there is a distinct community shift upon soil drying which is consistent with previous studies (Zhou et al. 2016; Pajares et al. 2018). Besides the confirmation that there are clear impacts on the community as a whole, we also noticed that new members of bacterial populations being more abundant in both conditions (dry and wet), which were not previously reported in a single study and/or similar soil profile(s). Our study thus provided the baseline that enabled each of the distinct bacterial populations to be studied further to enrich our current knowledge.

### 3.1. Soil drying influences the microbial community structure

Soil A generated ~64 million reads and soil B generated ~71 million reads. We assembled the reads with a minimum contig length of 1000bp, yielding ~47,000 and 51,000 contigs in soil A and B respectively (Supplementary Table S2). We used Kaiju ver 1.7.2 (Menzel, Ng, and Krogh 2016) to elucidate the impact of soil moisture content on the soil microbial community. Briefly, each of the reads in the metagenomes is assigned to a taxon in the NCBI BLAST non-redundant protein database. The reads are translated into amino acid sequences, and are searched in the database. We classified 365 families in both the soil locations (Supplementary Table S3). We showed a statistical difference (PERMANOVA, Pseudo-F 35.643, P < 0.05, Figure 1) in microbial community structure between the two distinct soil types (dry and wet).

**Figure 1:**
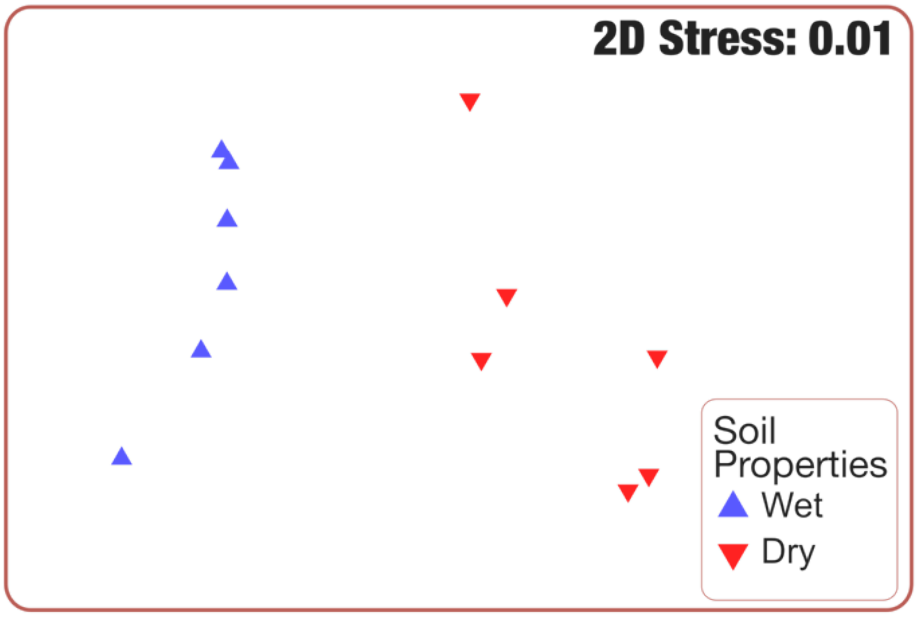
Microbial composition between wet (soil A) and dry (soil B) soil. Metagenomic shotgun sequencing data were assigned taxonomic identity using Kaiju and analyzed by Bray-Curtis dissimilarity between the two groups. NMDS plot shows that microbial composition from soil A and B formed distinct clusters.

We noticed, through post-hoc SIMPER analyses, that *Streptomycetaceae* (1.21%), *Bradyrhizobiaceae* (1.2%), *Vicinamibacteraceae* (1.13%), *Pseudonocardiaceae* (0.97%), *Planctomycetaceae* (0.9%) contributed the most in dry soil (Figure 2A, Supplementary Table S4). Following that, *Sphingomonadaceae, Burkholderiaceae, Hyphomicrobiaceae, Comamonadaceae, Conexibacteraceae, Mycobacteriaceae, Micromonosporaceae, Acidobacteriaceae, Rhizobiaceae, Rhodospirillaceae, Chitinophagaceae, Polyangiaceae, Methylobacteriaceae, Solibacteraceae, and Phyllobacteriaceae* also were among the top contributors. In the wet soil, *Bradyrhizobiaceae* (1.57%), *Streptomycetaceae* (1.21%), *Hyphomicrobiaceae* (0.99%), *Pseudonocardiaceae* (0.97%), and *Vicinamibacteraceae* were the top contributors (Figure 2B, Supplementary Table S5). *Micromonosporaceae*, *Chitinophagaceae*, *Acidobacteriaceae, Mycobacteriaceae, Burkholderiaceae, Conexibacteraceae, Sphingomonadaceae, Planctomycetaceae, Comamonadaceae, Solibacteraceae, Polyangiaceae, Nocardioidaceae, Methylobacteriaceae, Rhizobiaceae, Gemmatimonadaceae*, also contributed in wet soil.

**Figure 2:**
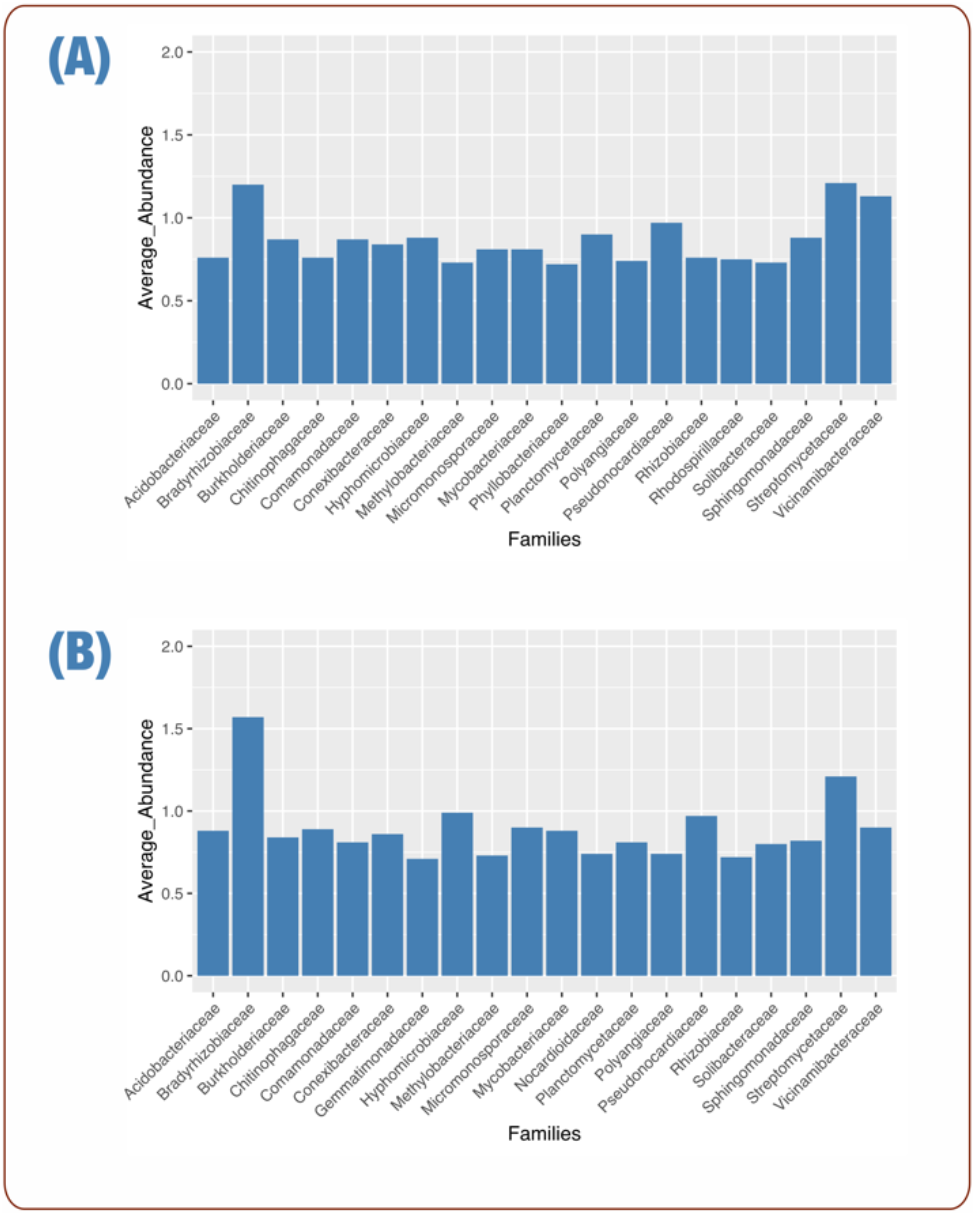
Key bacterial populations in wet and dry soil. Taxonomy of bacterial populations assigned using Kaiju showing (A) the relative abundance of *Streptomycetaceae, Bradyrhizobiaceae, Vicinamibacteraceae, Pseudonocardiaceae*, and *Planctomycetaceae in* dry soil. (B) the relative abundance of *Bradyrhizobiaceae, Streptomycetaceae, Hyphomicrobiaceae, Pseudonocardiaceae*, and *Vicinamibacteraceae* in wet soil.

*Vicinamibacteraceae* (dry: 1.13%, wet: 0.9%)*, Sinobacteraceae* (dry: 0.59%, wet: 0.4%)*, Xanthomonadaceae* (dry: 0.65%, wet: 0.52%)*, Christensenellaceae* (dry: 0.27%, wet: 0.16%)*, Planctomycetaceae* (dry: 0.9%, wet: 0.81%)*, Rhodospirillaceae* (dry: 0.75%, wet: 0.68%)*, Woeseiaceae* (dry: 0.27%, wet: 0.2%)*, Comamonadaceae* (dry: 0.87%, wet: 0.81%)*, Acidiferrobacteraceae* (dry: 0.36%, wet: 0.3%), were the top contributors that showed statistically higher average abundance in dry as compared to wet soil (Figure 3, Supplementary Table S6). On the other hand, the wet soil as compared to dry soil, showed statistically higher average abundance of *Bradyrhizobiaceae* (dry: 1.2%, wet: 1.57%), *Acidobacteriaceae* (dry: 0.76%, wet: 0.88%)*, Geodermatophilaceae* (dry: 0.45%, wet: 0.57%)*, Nocardioidaceae* (dry: 0.63%, wet: 0.74%)*, Hyphomicrobiaceae* (dry: 0.88%, wet: 0.99%)*, Micromonosporaceae* (dry: 0.81%, wet: 0.9%)*, Propionibacteriaceae* (dry: 0.43%, wet: 0.51%)*, Mycobacteriaceae* (dry: 0.81%, wet: 0.88%), among others. During our analysis of the top-contributors (Supplementary Table S6), *Bradyrhizobiaceae* (dry: 1.2%, wet: 1.57%) was one of the families that had high average abundance and contribution in both the wet and the dry soil, suggesting that soil moisture had little significant impact on its presence.

**Figure 3:**
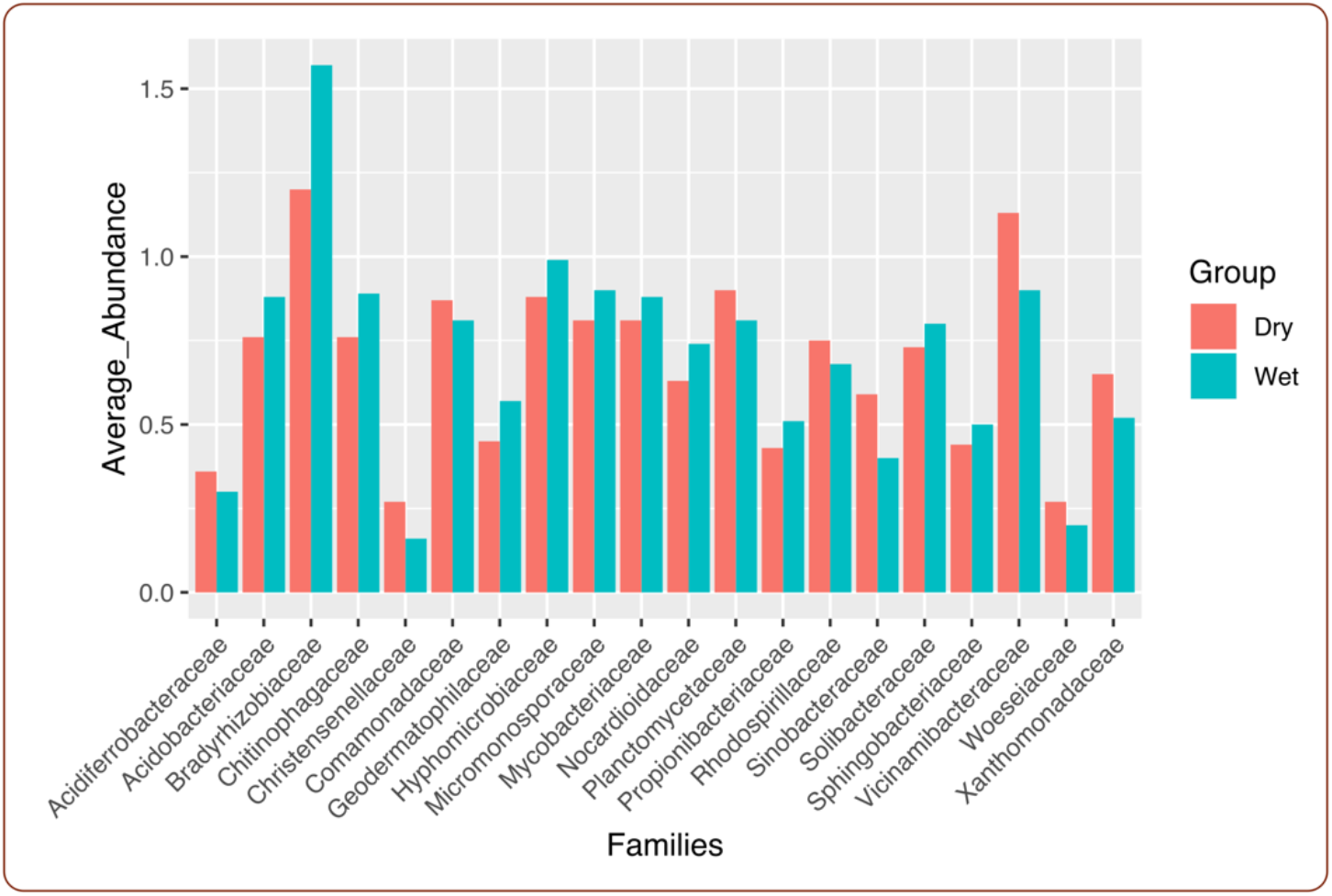
Top bacterial populations contributors for both dry and wet soil. Metagenomic shotgun sequencing data was analyzed using PERMANOVA and top contributors were identified using post-hoc SIMPER test.

Our metagenome analyses inferred that the soil moisture content dictated the microbial composition. In the previous studies, dry soil induced a similar microbial population profiling (Chowdhury et al. 2019). We further discovered that *Actinobacteria*, *Acidobacteria*, *Proteobacteria, Planctomycetes, Bacteroidetes* were the top contributing bacterial populations in soil with a lower moisture content, and may have significant biological importance under drought conditions. For example, *Actinobacterium* isolated from semi-arid environments has been reported to enhance the drought tolerance in maize (Selim et al. 2019), while *Acidobacteria, Proteobacteria, and Bacteroidetes* are ubiquitous in soils thus explaining its importance (Chowdhury et al. 2019; Chodak et al. 2015). *Acidobacteria* has been reported to exist in nutrient poor soil with low carbon availability (Fierer, Bradford, and Jackson 2007). *Planctomycetes*, on the other hand, has some interesting features: (a) large genomes (b) lack of peptidoglycan in the cell walls, and (c) division of cells by inner membranes into separate compartments, which might be responsible for its contribution and resistance towards drought (Daniel H. Buckley et al. 2006).

SIMPER analysis indicated that there were ~110 families that were overlapping in terms of average relative abundance in both wet and dry soil (Supplementary Table S7). In order to understand the true picture of the community shift, we looked into the families that were exclusive to the dry and the wet. The bacterial families that were found to be present exclusively in the wet soil include *Akkermansiaceae, Beijerinckiaceae*, and *Glycomycetaceae. Akkermansiaceae* belongs to the phylum *Verrumicrobia*. Members of the phylum *Verrumicrobia* are ubiquitous in soil, and are extremely sensitive to environmental alterations, which might explain its presence in wet and not in dry soil in this study (Kielak et al. 2008; Bruce et al. 2010; D. H. Buckley and Schmidt 2001; Pan et al. 2014; Acacio Aparecido Navarrete, Diniz, et al. 2015; Acacio A. Navarrete et al. 2015; Acacio Aparecido Navarrete, Soares, et al. 2015). *Beijerinckiaceae* shows an ability of fixing nitrogen, even able to grow in nitrogen limiting conditions (Marín and Arahal 2014), providing essential conditions in aiding plant growth (Thuler et al. 2003; Miyasaka et al. 2003). *Glycomycetaceae* are soil isolates, belonging to the phylum Actinobacteria (Stackebrandt 2014).

On the other hand, we identified *Alicyclobacillaceae, Christensenellaceae, Oceanospirillaceae, Rhodocyclaceae, Vulgatibacteraceae, and Woeseiaceae*, which were exclusively in dry soil. Members of the *Alicyclobacillaceae* are capable of forming endospores and have been isolated from extreme environments, and it might explain why this family was thriving in dry soil conditions (Imperio, Viti, and Marri 2008; Goto et al. 2003). *Christensenellaceae* are well-known for their characteristics of being fermentative, and demonstrating the capabilities of reducing iron and sulfur, which might allow them to show its existence in such a water stressed environment (Gupta et al. 2018). *Oceanospirillaceae* has been previously isolated from soils contaminated with oil-sludge and oil-pollutions. So, this supports the fact of its prevalence in the drier soil here (Kuang et al. 2018; Chikere et al. 2019). Members of the *Rhodocyclaceae* have been isolated from a diverse range of environments i.e. soil, plants dealing with sewage treatments, polluted and unpolluted water sources, plant roots (Oren 2014). Their existence in an extreme soil environment might be due to their properties of being able to utilize a range of carbon sources like nitrate, oxygen, selenate, chlorate, and perchlorate (Oren 2014). They are primarily anoxygenic photoheterotrophs, and can even function as nitrogen-fixing aerobes, sulfur-oxidizing, methylotrophic, and propionic-acid fermentation (Oren 2014). *Vulgatibacteraceae* has been previously isolated from forest soils (Yamamoto, Muramatsu, and Nagai 2014), and *Woeseiaceae* from coastal sediments (Coskun et al. 2019). *Woeseiaceae* are carbon-fixing microbes which oxidizes compounds with reduced sulfur in extreme coastal environments (Coskun et al. 2019).

However, associating direct functional mechanisms to such a diverse group(s) can be misleading as there are so many species and strains within each phylum and family. We are motivated to conduct further functional analyses at the species level, in order to elucidate the genomic functions that are essential to bacterial populations during drought conditions.

### 3.2. Soil moisture has a significant impact on selected bacterial genomes and function

We used post-hoc SIMPER analyses to detect the contribution to dissimilarity at the species level between the dry and wet soil, and used that information to select fifteen bacterial genomes for further gene level analyses (Figure 4, Supplementary Table: S8).

**Figure 4:**
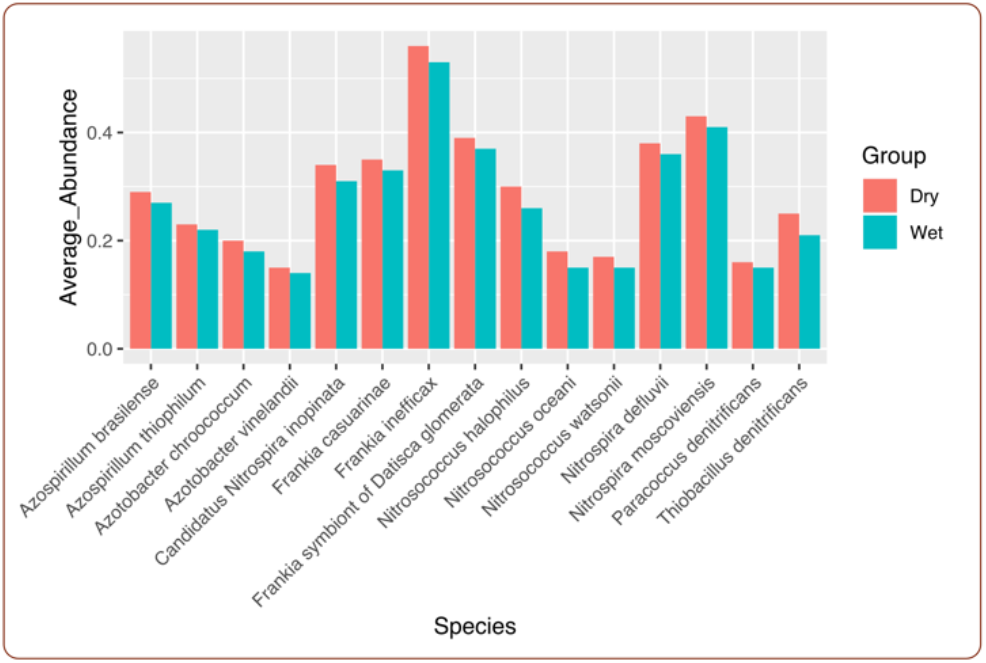
Distribution of the 15 downloaded key genomes in wet and dry soil. Relative abundance of the 15 genomes in the dry and wet soil, showing that all the 15 genomes have a higher relative abundance in the dry soil than in wet. *Nitrosococcus halophilus* and *Thiobacillus denitrificans* show the highest relative abundance in the dry soil when compared to wet among the 15 genomes that we selected.

We used Bowtie2 (Langmead and Salzberg 2012) to map the 12 metagenomes to the 15 downloaded bacterial genomes. We performed PERMANOVA statistical analyses, and showed that soil drying had a significant effect on these bacterial populations (PERMANOVA, Pseudo-F 7.2352, P < 0.05, Figure 5). We observed that *Azotobacter vinelandii* (dry: 1.82%, wet: 1.51%)*, Azotobacter chroococcum* (dry: 1.96%, wet: 1.66%)*, Nitrosococcus watsonii* (dry: 2.04%, wet: 1.75%)*, Nitrosococcus halophilus* (dry: 1.8%, wet: 1.52%)*, Nitrosococcus oceani* (dry: 1.92%, wet: 1.66%)*, and Thiobacillus denitrificans* (dry: 2.84%, wet: 2.67%) had a higher average relative abundance in dry soil than in wet (Supplementary Table: S9). On the other hand, *Frankia casuarinae* (dry: 3.93%, wet: 4.34%)*, Frankia inefficax* (dry: 3.11%, wet: 3.51%)*, Frankia Datisca glomerata* (dry: 3.86%, wet: 4.15%), showed a higher average relative abundance in wet soil (Supplementary Table: S9).

**Figure 5:**
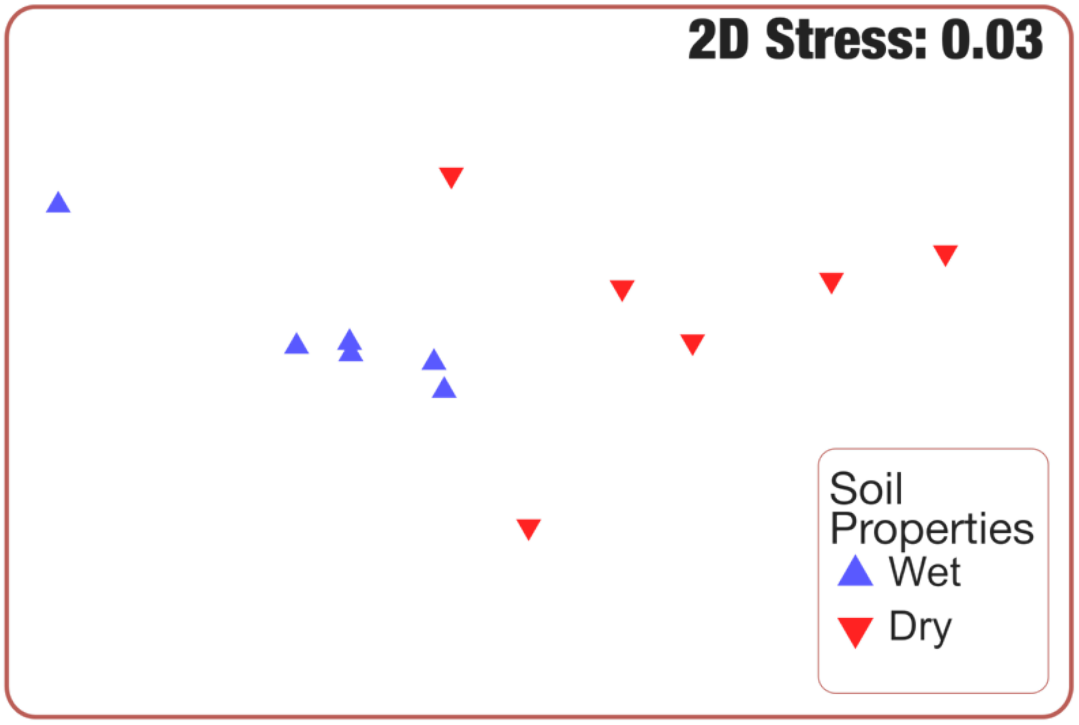
Relative abundance of 15 downloaded key genomes in wet and dry soil. Metagenomic shotgun sequencing data were mapped to the 15 downloaded genomes and analyzed by Bray-Curtis dissimilarity between the two groups. NMDS plot shows that the relative abundance of the 15 genomes composition from soil A and B formed distinct clusters.

It was evident that the genus *Azotobacter*, *Nitrosococcus*, and *Thiobacillus* were more prevalent in dry conditions, and *Frankia* was more abundant in wet soil. We believed that being the fact that it is a very well-known biological nitrogen-fixer (Santi, Bogusz, and Franche 2013) was one of the reasons *Frankia* is present in both wet and dry soil. Interestingly, *Azotobacter* is capable of surviving in dry soil conditions, resisting tough environmental conditions such as drying or radiations (Vela 1974; Socolofsky and Wyss 1961; Velva and Wyss 1965), by showing some dormant structures, reported to be cysts (Socolofsky and Wyss 1961). These reports corroborate well with our studies that find them more abundant in dry soil. The higher average relative abundance of *Thiobacillus denitrificans*, *Nitrosococcus watsonii, Nitrosococcus halophilus, Nitrosococcus oceani* in the dry soil can be potentially interesting as these species are related to various biogeochemical cycles (Prosser 2005; Pajares and Bohannan 2016). Based on these results and subsequent analyses, we hypothesized that these selected bacterial genomes on a whole were influenced by soil moisture content, and that we exploited genomics information within these genomes to gain further insights into their mechanisms during times of drought stress.

We used post-hoc SIMPER analysis on all of the genes in each of the 15 downloaded genomes (Supplementary Table S10). SIMPER test yielded gene lists of the top genes with average relative abundance in both types of soils. Based on that, we selected ~top 20 genes in most of the 15 downloaded genomes (Supplementary Table S10). Among those genes, we observed a higher coverage of translation elongation factor EF-Tu, thiamine biosynthesis protein, and catalase among the fifteen genomes in which these genes were present in the dry as compared to the wet soil, and speculated that these three genes are crucial in enabling bacterial populations to be more resilient during drought and/or high heat conditions (Figure 6).

**Figure 6:**
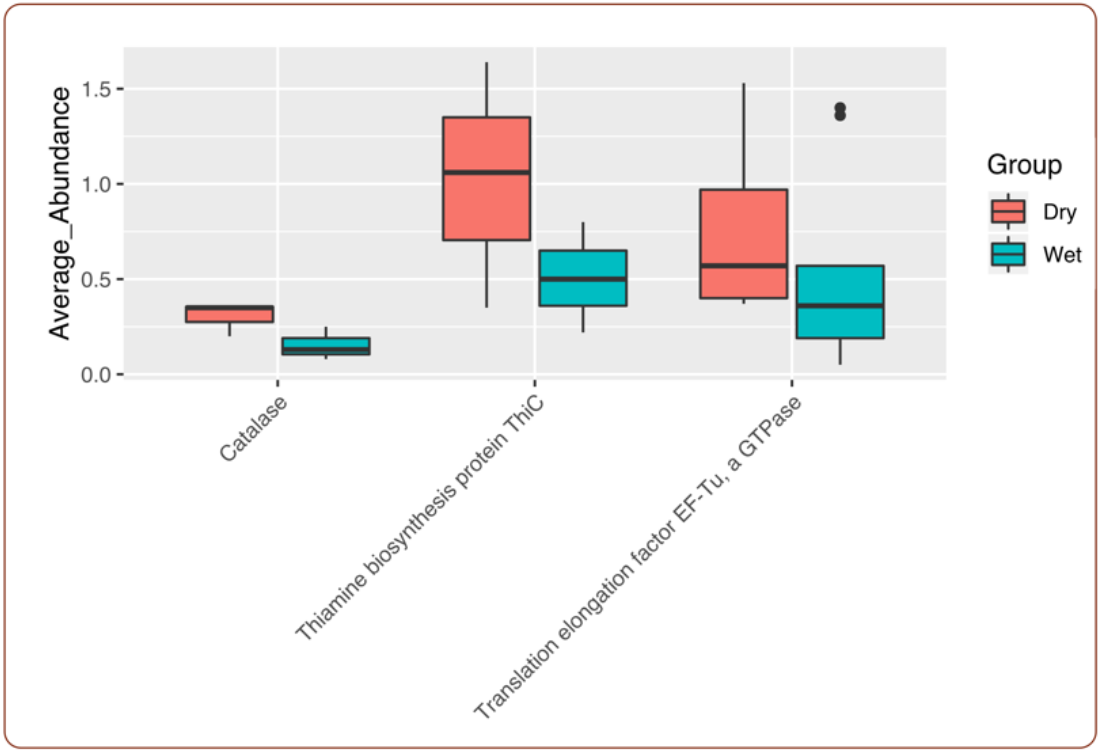
Distribution of EF-Tu, thiamine biosynthesis, and catalase gene function in wet and dry soil. EF-Tu, thiamine biosynthesis, and catalase within the 15 genomes have significantly higher relative abundance in dry as compared to wet soil. These three bacterial genes are associated with microbial resilience during drought conditions.

We noticed that translation elongation factor thermo unstable (EF-Tu) was one of the top-contributing genes in *Azosprillum brasilense, Azospirillum thiophilum, Azotobacter vinelandii, Candidatus Nitrospira inopinata, Frankia casuarinae, Frankia inefficax, Frankia Datisca glomerata, Nitrospira moscoviensis*, and *Paracoccus denitrificans* (Figure 6, Supplementary Table S11), suggesting that EF-Tu might be one of the mechanisms in which bacterial populations used in overcoming stressful conditions. EF-Tu is one of the most abundant proteins that is found in the bacteria and contributes to almost 6% of the total expressible protein in *Escherichia coli (Furano 1975)*. Although the primary role of EF-Tu is attaching aminoacyl-tRNA to the ribosome of the cells (Sprinzl 1994), there are other significant diverse roles in various microorganisms. In bacteria, EF-Tu has also been found to protect proteins from stress by forming stable complexes under heat shock conditions, and thus aiding in protein folding and renaturation in the cytoplasm (Caldas, El Yaagoubi, and Richarme 1998) Thus, there have been reports for EF-Tu to protect proteins during heat stress, and assist folding of a number of proteins such as citrate synthase and α-glucosidase (Caldas, El Yaagoubi, and Richarme 1998).

We also noticed in our study, thiamine biosynthesis protein was one of the top-most contributing genes in *Azotobacter chroococcum, Azotobacter vinelandii, Frankia Datisca glomerata* (Figure 6, Supplementary Table S12). Our data suggested that these bacterial populations may use thiamine biosynthesis protein to synthesize thiamine, helping these bacterial populations to respond directly to the drought induced oxidative stress by interacting with free radicals (Wolak et al. 2014). In a more indirect approach, synthesized thiamine can also function as a cofactor as transketolase and α-ketoglutarate dehydrogenase which influences the redox status of the cells (Bunik 2003; Rapala-Kozik, Kowalska, and Ostrowska 2008). In *Escherichia coli*, there are reports that thiamine triphosphate (TTP) and its derivative (AThTP) overexpressed under amino acid or carbon stress, acting as ‘alarmones’ (Lakaye et al. 2004; Gigliobianco et al. 2010).

Similarly, our analysis showed catalase contributed most in *Azospirillum brasilense, Frankia casuarinae*, and *Frankia Datisca glomerata* (Figure 6, Supplementary Table S13). We hypothesized that catalase, being a popular antioxidant enzyme known for detoxification of reactive oxygen species (Schellhorn 1995), enabled the bacterial populations to survive stress by degrading molecules such as H_2_O_2_. Catalase is omnipresent in a range of species from bacteria to humans, and neutralizes the toxic H_2_O_2_, converting to oxygen and water (Zamocky, Furtmüller, and Obinger 2008). Currently, there are three classes of catalase, namely: catalase, catalase-peroxidase, and pseudocatalase (X. Liu and Kokare 2017), each class involved in performing the reaction: (2 H_2_O_2_ → 2 H_2_O + O_2_). We proposed that our short-listed genomes should be divided into three distinct groups and a common mechanism that might be contributing to each group’s functioning and resilience to dry conditions (Supplementary Figure 1). In our study, EF-Tu, thiamine biosynthesis, and catalase had more coverage in dry soil than in wet in each of the genomes, and also belong to the top contributing genes in the respective organisms. Our results suggested that bacterial populations may use these mechanisms to enhance their resistance in terms of drought stress.

Our study summarizes the shifts in the microbial structure, as well as in the functions impacted by soil dryness. There were definite impacts of soil dryness on the bacterial community. In dry soil, we demonstrated that there was an increase in the average relative abundance of some families and species, which might have important biological significance(s). Moreover, by short-listing some important bacterial populations, we explored their potential pathways, and deciphered genes such as EF-Tu, thiamine biosynthesis, and catalase can be key to those microorganisms in aiding their resistance against drought stress. To conclude, in this study, we not only aimed at elucidating the shifts in the bacterial population at the community level and at the genome level but also explore further into the potential mechanisms that the population might use in moisture-deprived environments. This research approach can be applied globally and across different environmental systems, providing insights into deciphering the bacterial community composition and the microbial functional mechanisms.

## 4. Acknowledgments

Soumyadev Sarkar acknowledges the National Science Foundation EPSCoR for his research grant. Kaitlyn Ward was supported by the research award from the College of Arts and Sciences, Kansas State University.

## Conflict of interests

All authors declare that they have no conflict of interest.

---

Supplementary Table 1: Details of the 15 downloaded genomes.

Supplementary Table 2: Assembly statistics.

Supplementary Table 3: Classification of 365 families in both the soil locations.

Supplementary Table 4: Families that were top contributors in the dry soil.

Supplementary Table 5: Families that were top contributors in the wet soil.

Supplementary Table 6: SIMPER analysis indicated the top contributors in dry soil when compared to wet.

Supplementary Table 7: SIMPER analysis detected ~110 families that were overlapping in terms of average relative abundance in both wet and dry soil. Families that were exclusive to the dry and the wet are indicated with red colored font.

Supplementary Table 8: Relative abundance of 15 downloaded genomes in dry and wet soil.

Supplementary Table 9: Bacterial population among 15 genomes that show difference in relative abundance and also belong to the top contributors.

Supplementary Table 10: SIMPER analysis on all of the genes in each of the 15 downloaded genomes and ~top 20 genes in most of the 15 downloaded genomes.

Supplementary Table 11: Distribution of EF-Tu gene function in wet and dry soil.

Supplementary Table 12: Distribution of thiamine biosynthesis gene function in wet and dry soil.

Supplementary Table 13: Distribution catalase gene function in wet and dry soil.

Supplementary Figure 1: Individual pathways for the genomes.

## Notes

### Competing Interest Statement

The authors have declared no competing interest.

